# Visual cue properties determine innate orientation strategy in Monarch butterflies

**DOI:** 10.64898/2026.06.16.732693

**Authors:** Fredrik Øglænd Hanslin, Maria Gayler, Myriam Franzke, Basil el Jundi

**Affiliations:** Department of Biology, Norwegian University of Science and Technology, Trondheim, Norway; Department of Zoology II, Biocenter, University of Würzburg, Germany; Institute of Biology and Environmental Sciences, University of Oldenburg, Germany

**Keywords:** insect, *Danaus plexippus*, lepidoptera, landmark, stripe fixation, attraction behavior, compass, menotaxis, navigation

## Abstract

Animals rely on a wide range of environmental signals, including celestial and terrestrial cues for navigation. While celestial cues, such as the sun, play a major role in maintaining a constant heading during long-distance migration and dispersal, terrestrial cues provide an animal with a short-range navigation system, ideal to pinpoint highly specific locations. In Monarch butterflies, the simulation of a terrestrial landmark, i.e. a vertical stripe, induces an attraction behavior (all animals head toward the stimulus) while a small green light spot, simulating the sun, elicits menotactic orientation (animals adopt individual-specific headings relative to the stimulus). However, the mechanisms underlying how the animal distinguishes between a stimulus as a terrestrial landmark versus a celestial cue remains unclear. To explore this, we tested non-migratory Monarch butterflies (*Danaus plexippus*) in a flight simulator. The inner surface of simulator was equipped with an area of LEDs, allowing to present different visual stimuli to the butterflies during tethered flight. By systematically manipulating the stimulus’ width, height, brightness, and elevation we found that Monarch butterflies exhibited attraction behavior to high contrast areas, like stripe edges. Menotactic behavior was not achieved by solely decreasing the stimulus to a small light spot but also required for the stimulus to be presented at higher elevation to be interpreted as a sun stimulus. These findings suggest that multiple parameters, inherently set by the butterfly’s navigation system, are critical to interpret a visual stimulus as celestial cue or terrestrial landmark, producing dynamic switches between different orientation strategies during navigation.

## INTRODUCTION

To produce a variety of behaviors, insects must constantly extract relevant sensory information from their environment to keep track of their movements through space. In this context, global cues, such as the position of the sun and other celestial cues (Dreyer et al., 2025; el Jundi et al., 2014; Freas and Spetch, 2023) but also terrestrial visual landmarks play an important role as orientation references. Terrestrial visual landmarks are especially crucial for short-range navigation enabling insects to find specific locations (de Ibarra et al., 2015; Richter et al., 2023). These visual landmarks support different orientation and navigation strategies (Dill et al., 1995; Maimon et al., 2008; Ofstad et al., 2011). For instance, bees (Dittmar et al., 2011; Lehrer and Collett, 1994), ants (Buehlmann et al., 2020; Collett et al., 1992), and flies (Ofstad et al., 2011) use visual landmarks to memorize the location of biologically relevant sites. In other contexts, insects rely on visual landmarks to be attracted to a specific spot or to maintain a constant bearing relative to the visual cue (Collett, 1988; Franzke et al., 2022). Interestingly, laboratory experiments on many insects have shown that specific features of a visual landmark, such as shape, size, and orientation, strongly influence the orientation behavior (Jander, 1971; Liu et al., 2006). For example, flies are attracted by a longer vertical stripe stimulus, which they seem to interpret as a landing site, but are repelled by a shorter stripe, which may be perceived as a predator (Maimon et al., 2008; Mongeau et al., 2019). While recent work on looming sensitivity suggests that stimulus interpretation arises from dynamic interactions between sensory and motor circuits in the brain (Ache et al., 2019), the precise mechanisms underlying context-dependent orientation strategies in insects remain poorly understood.

Monarch butterflies (*Danaus plexippus*) exhibit two distinct phenotypes, a migratory and a non-migratory one. The migratory phenotype is renowned for its annual long-distance migration from North America to Central Mexico (Mouritsen, 2018; Reppert and de Roode, 2018). During this journey, the butterflies primarily rely on a time-compensated sun compass to maintain their migratory heading (Merlin et al., 2009; Mouritsen and Frost, 2002; Perez et al., 1997; Sauman et al., 2005) though they may also rely on the polarization pattern of the sky (Reppert et al., 2004; Sauman et al., 2005, but see also Stalleicken et al., 2005) and other skylight cues for orientation (el Jundi and Warrant, 2026; Goforth and Merlin, 2025; Grob et al., 2025). The non-migratory phenotype typically disperses locally over short distances (Fisher et al., 2020). The most effective dispersal strategy involves maintaining a straight flight heading in a random direction, a behavior known as menotaxis (Grob et al., 2021). During dispersal, Monarch butterflies likely combine sun-related cues with terrestrial visual landmarks such as the panoramic skyline, to keep and stabilize their heading (Franzke et al., 2020). Furthermore, they need to rely on precise short-range navigation to find food sources and locate suitable habitats for egg-laying and mating sites (Fisher and Bradbury, 2021; Grob et al., 2025; Konnerth et al., 2023). Taken together, the combination of sun-related cues and terrestrial visual information seem to play an important role for the Monarch butterfly heading system, allowing the butterflies to navigate over large distance as migrants or find suitable habitats as non-migrants.

Like flies, non-migratory Monarch butterflies dynamically adjust their orientation strategy based on the prevailing visual scene. Thus, a long vertical stripe elicits attraction behavior, whereas a small, bright spot, simulating the sun, induces menotactic flight behavior (Franzke et al., 2022). However, the exact sensory and behavioral mechanisms that determine whether an insect relies on menotaxis or phototaxis/attraction remains unclear. Here, we conducted well-controlled flight simulator experiments within an LED arena to define the sensory and behavioral rules that underlie these different orientation strategies. We systematically manipulated key features of the visual scene, such as width, height, brightness, and elevation of a stripe to investigate how non-migratory Monarch butterflies responded to changes in their visual world. Crucially, we asked under which visual setting butterflies “interpret” a visual stimulus as “sun” (eliciting menotactic behavior) versus a terrestrial landmark (eliciting attraction). This approach allowed us to identify the perceptual filters that govern context-dependent orientation in Monarch butterflies.

## MATERIALS AND METHODS

### Animals and Preparation

We tested non-migratory Monarch butterflies (*Danaus plexippus*), which were ordered as pupae from Costa Rica (Costa Rica Entomology Supply, butterflyfarm.co.cr). They were kept in an incubator (HPP 110 und HPP 749, Memmert GmbH+Co. KG, Schwabach, Germany) at 25 °C, 80% relative humidity and a 12:12h light/dark cycle. After eclosion, the butterflies were individually marked and transferred into a different incubator (I-30VL, Percival Scientific, Perry, IA, USA) at 50% relative humidity and a light-dark cycle of 12:12h. In the light phase, the temperature was set to 25 °C while it was adjusted to 23 °C in the dark phase. The adult butterflies had ad libitum access to feeders filled with a 15% sucrose solution.

Prior to testing, we prepared the butterflies by removing the scales from their dorsal thorax and gluing (EVO-STIK, Bostik Ltd, Stafford, UK) a tungsten stalk to it (0.508×152.4 mm, Science Products GmbH, Hofheim, Germany). To allow the glue to harden, the butterflies were subsequently kept in plastic cups in darkness for around three hours with ad libitum access to 15% sucrose solution.

### Experimental Setup and Procedure

To explore how Monarch butterflies use different visual cues, we conducted a set of experiments indoors in a cylindrical LED-flight simulator (Fig. 1A). For each trial, one animal was tethered in the middle of the arena with the tungsten stalk connected to an optical encoder (E4T miniature Optical Kit Encoder, US Digital, Vancouver, WA, USA). The encoder recorded the heading directions every 200 ms with an angular resolution of 3°. The data were sent to a computer via a digitizer (USB4 Encoder Data Acquisition USB Device, US Digital, Vancouver, WA, USA) running the corresponding software. To be able to present different visual stimuli to a butterfly, the inner surface of the flight simulator was equipped with RGB-LED-panels with 2,048 LEDs in 16 rows and 128 columns (16×16 APA102C LED Matrix, iPixel LED Light Co., Ltd, Baoan Shenzhen, China). The LEDs were controlled via a Raspberry Pi Computer (Raspberry Pi 3 Model B, Raspberry Pi Foundation, UK).

**Fig. 1.**
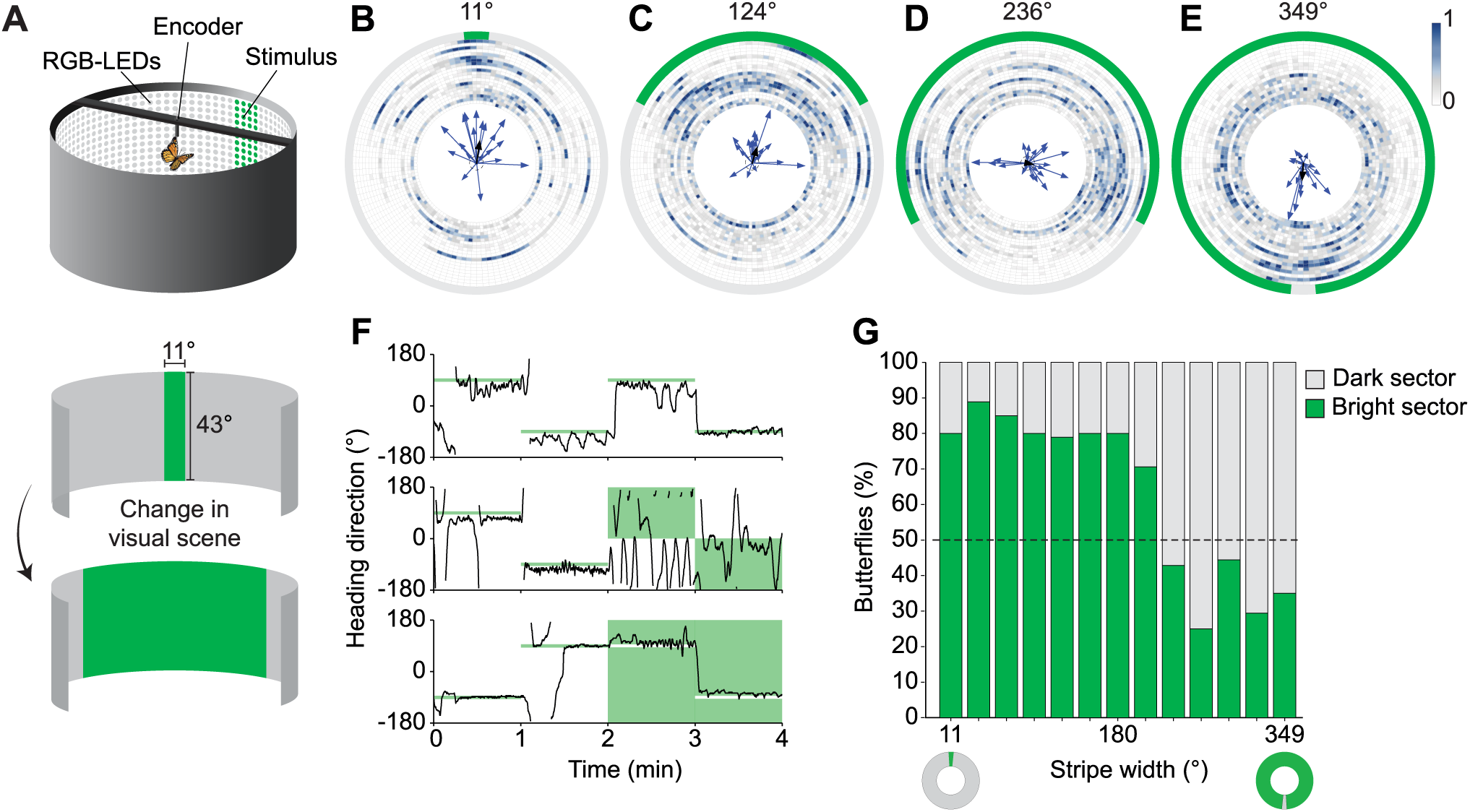
Landmark orientation to a vertical stripe of varying width. (A) Schematic of the experimental setup displaying the flight simulator equipped with 2048 RGB LEDs (*top*). Butterflies were tethered to an optical encoder that recorded the heading directions when presented with the stripe stimulus of varying width (*bottom*). (B, C, D, E) Mean heading directions and vector strengths displayed inside circular heatmaps of raw heading directions for four different experimental conditions, with stripe sizes shown as the green perimeter. The heatmaps display the normalized count of heading directions, with each trial split into separate rings, sorted by increasing vector strength starting from the center. (F) Heading direction of exemplary individuals for three different conditions (*top*: green stripe width was set to 11° throughout the entire experiment; *middle*: green stripe width changed from 11° to 180° after two minutes; *bottom*: green stripe width changed from 11° to 349° after two minutes). The position of the visual cue was displaced by 180° every minute. Green shaded area indicates the stripe width and condition. (G) Bar plot of the proportion of butterflies that display mean directions towards the dark or bright sector of the flight simulator for the different stripe widths.

### Variable stripe width

In the first set of experiments, we studied how the butterflies changed their orientation behavior with respect to a vertical green stripe (vertical extend: 43°, emission peak at 520 nm) that varied in widths between 11° and 349°. The orientation of every butterfly was recorded for a total of four minutes. In the first two minutes we presented a narrow stripe (11°) to the butterflies. This stripe changed its position by 180° in the flight arena after one minute, to ensure that the butterfly was actively orienting to the cue. Thus, animals that used the stripe to maintain a constant flight direction were expected to change their heading by about 180°. In the second two minutes, we presented either the same stripe width (*control*) or a wider vertical stripe (*test*) to the butterflies (n = 20). In the experiments with a wider stripe, the stripe width was adjusted to either 39° (n = 18), 68° (n = 20), 96° (n = 20), 124° (n = 19), 152° (n = 20), 180° (n = 19), 208° (n = 17), 236° (n = 21), 264° (n = 20), 293° (n = 18), 321° (n = 17) or 349° (n = 18). Again, the position of the stimulus was changed by 180° after each minute to test if the butterflies followed the relocation of the stripe. Thus, each butterfly experienced in total three turns of the visual stimulus. Half of the animals started with the stripe at an angular position of 90° (relative to the orientation of antero-posterior body axis of the animals at the beginning of the experiments), the other half with the stripe presented at 270°. The intensity measured in the middle of the arena facing towards the center of the stripe ranged from 2.43×10^13^ photons/cm^2^/s (stripe width: 11°) to 1.85×10^14^ photons/cm^2^/s (stripe width: 349°).

### Vertical stripe height

In the second set of experiments, we tested the influence of changes in the stripe’s vertical height (stripe width: 11°) on the butterflies’ orientation over six minutes. The center of the stripe was always at 0° elevation relative to the butterflies’ eye level. In the first two minutes, we again presented a long stripe (43° high) as a control to the butterflies, before presenting stripes of three different vertical extends (22°, 11° and 5°) as well as a phase with no stripe, in randomized orders. After each minute, the stimulus was turned by 180° to ensure that the butterfly was actively orientating to the cue. This experiment was conducted with a green stripe on a dark background (n = 19) as well as with a dark stripe on a green background (n = 18).

To test the effect of cue elevation on the butterflies’ orientation strategy, we repeated the previous experiment with the green stripe, now gradually increasing the elevation of the cue as the vertical extent become shorter (n = 20). Thus, the butterflies were first presented with a long stripe, before presenting stripes of three different vertical extends (22°, 11°, and 5°), with respective elevations (11°, 17°, 19°) at the stripe center, in addition to a phase with no stripe. Similar to the previous experiment, the order of the different visual stimuli was randomized.

## Data analysis

We analyzed our data with the software MATLAB (R2018a, Version 9.4.0.813654 MathWorks, Natick, MA, USA) and the CircStat toolbox, and R (version 4.3.2) with a significance level of a = 0.05. Each trial was divided into 1-min phases because the stimulus was turned by 180° every minute, to ensure that the animal was actively orientating.

To test whether the animals followed the turning of a stripe, we calculated their change of heading as the angular difference of the mean directions between two consecutive phases. For each animal, a mean vector within every phase was calculated showing its mean heading direction as well as its vector strength (r), a measure of directedness (Batschelet, 1981). The vector strength ranges from 0 (completely disoriented) to 1 (perfectly oriented in one direction).

As the results of our experiments can be affected by the animals’ context and motivation, we had to exclude some animals from our analysis (variable width: n = 243, variable height: n = 80). It is important to note that the following filtering criteria were set prior to conducting all experiments and were not modified afterwards. In all experiments with a bright cue on a dark background, animals with a vector strength < 0.1169 in any of the phases were excluded from the analysis. This value is shown to be the threshold for directedness in Monarch butterflies when presented with a cue that contains no directional information (Franzke et al., 2020). We favor the vector strength threshold over the Rayleigh test for non-uniformity when looking at individual trials, as the individual mean vectors are calculated over a very high sample size, resulting in a larger proportion of false positives when using the Rayleigh test (Rayleigh, 1880). In the experiments with the variable height of the stripes, we had to study the orientation behavior of Monarchs in darkness, which is not trivial given that they in general stop flying when no cue is presented in our indoor flight simulator. In this condition, we used a less conservative filter (vector strength > 0.2) similar to what has been used in previous experiments (Dreyer et al., 2021).

This threshold was chosen because butterflies that do not have any visual cue available in a flight simulator should never show a vector strength (r) exceeding 0.2 (Franzke et al., 2022). Those animals likely relied on non-visual cues, such as idiothetic cues (Beetz et al., 2022; Kraus et al., 2026) to keep their constant flight heading in darkness. In addition, we also excluded all butterflies that did not turn by 180° ± 90° with the relocated stimulus in the control phases at the beginning of every experiment. This was done, as the subsequent experimental design was based on the assumption that the butterflies relied on the presented visual cue for orientation, instead of idiothetic cues or any latent non-visual cues.

For simplicity, we presented all heading data relative to the azimuth of the center of the stripe (0°). For the experiments regarding the width of a stripe, the second two minutes were treated as one 2-minute phase. The randomized phases of the variable stripe height experiments were ordered according to the stripe height, starting with the longest stripe, and decreasing until no cue was left. For every group we used the non-parametric Moore’s modified Rayleigh test (MMRT) to test for a preferred population heading direction during each phase, and thereafter calculated the weighted mean direction, vector strength, and 95% confidence interval (Moore, 1980).

In order to test if the butterflies followed the stripe edges, we first converted the raw flight directions (bin size = 10 s, N = 2964, n = 247), and the angular edge positions [-180°,180°] to absolute values [0°,180°]. In this transformation, the two edges of the corresponding stripe will be at the same angle, allowing us to compare the edges simultaneously despite potential bimodality in the heading directions. With these data, we conducted a linear quantile mixed model from the ‘lqmm’ package on the median (τ = 0.5) to handle the non-normally distributed response variable and included the individual trials as a random effect to account for the non-independence of data points within a trial.

To further investigate whether individual butterflies switched between the two edges, we defined each edge area as the stripe width v) centered at each edge, yielding two areas: 0, vv and -v, 0v. When the stripe width exceeded 180°, the stripe width was defined as 360° - v, with the stripe center at 180°, yielding two edge areas: 360° - v, 180°v and 180°, 360° + vv (Fig. 2C). From here, we could calculate the proportion 0,1v of time spent flying towards each edge area for each individual to study whether individuals fixate on one of the edges (proportion = 0, or proportion = 1), or spend equal amount of time in each area (proportion = 0.5).

**Fig. 2.**
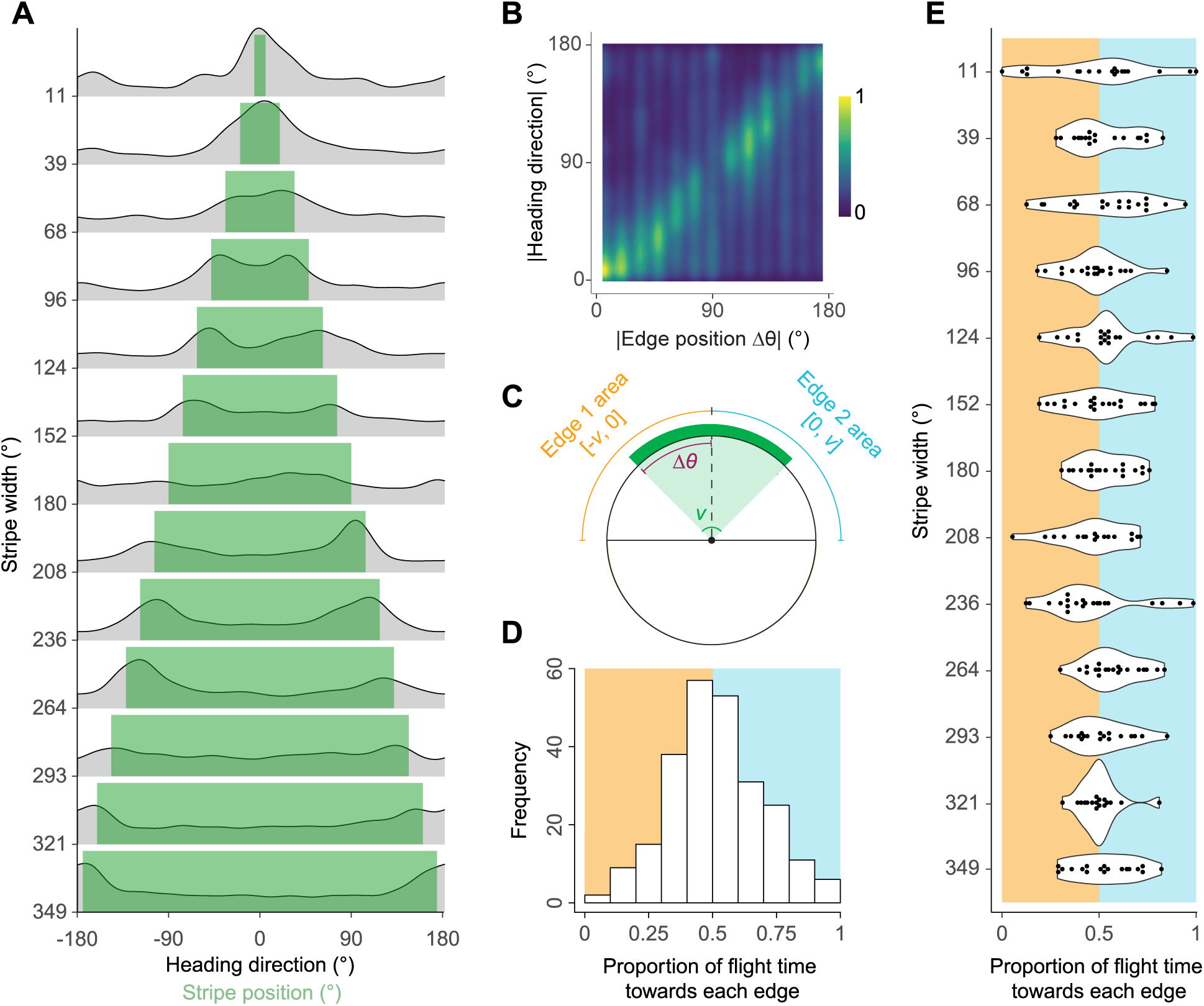
Flight performance relative to stripe edges. (A) Ridged density plots of the distribution of raw heading directions for each condition showing that heading directions are concentrated towards the stripe edges. Green area shows the stripe width and position of the edges. (B) Heatmap of the normalized absolute heading directions against the absolute stripe edge position, showing the strong relationship between heading direction and the edges. (C) Illustration to show the definitions of edge areas, where each area is explained by the stripe width (*v*) centered around each edge position (Δθ). (D) Histogram of proportion of flight time spent towards each edge area for each individual as defined in (C) across all conditions and (E) for each stripe width.

In addition to the Moore’s modified Rayleigh test, we also conducted inferential statistics to compare the orientation strategies between the centered stripe and elevated stripe experiments. We defined a binary variable a E 0,11 based on the individual mean heading direction, whether it was flying towards the cue (0° ± 90°, a= 1) or away from the cue (180° ± 90°, a = 0). We then implemented this variable as our response in a binomial Generalized Linear Model (GLM) with a logit link function using the ‘glmmTMB’ package, explained by the predictor variables stripe size, and experiment type (Centered, Elevated) in an interaction. Every model fit was validated using the ‘DHARMa’ package when applicable.

## Results

### The butterflies’ heading direction depends on stripe widths

In previous experiments, it has been established that non-migratory Monarch butterflies maintain constant flight headings with respect to visual cues (Franzke et al., 2020), based on different orientation strategies (e.g. menotaxis, phototaxis; Franzke et al., 2022). For instance, a stripe elicited an attraction behavior in Monarch butterflies with almost all animals keeping a directed course towards the stripe (Franzke et al., 2022). We now asked, if this attraction to the stripe is stable, irrespective of stimulus size. To study this, we tethered individual Monarch butterflies at the center of a flight simulator. The inner surface of the simulator was equipped with an array of LEDs, covering 360° around the animal (Fig. 1A). This allowed us to vary the azimuthal extent of a stripe, i.e., the width of the stimulus, while we recorded the heading direction of individual butterflies at the center of the simulator. As expected, when presented with a narrow, green stripe (11°, λ = 520 nm, intensity = 2.43×10^13^ photons/cm^2^/s), the butterflies maintained a directed flight towards the stripe (n = 20, p = 0.001, MMRT, 8 = 8°, 95% CI [328 - 48°], Fig. 1B, F top panel). This is consistent with what has been previously shown in Monarch butterflies when presented with a narrow green stripe (Franzke et al., 2022). As the width of the cue increased, we observed larger variations in the individual heading directions, still concentrated towards the center of the cue (n = 19, p = 0.003, MMRT, 8 = 17°, CI [320 - 74°], Fig. 1C, S1A). However, as the stripe’s width continued to increase beyond 180°, the heading directions started to stray away from the cue center, concentrating more around the stripe edges, resulting in a non-directed population (n = 21, p = 0.738, MMRT, Fig. 1D). Interestingly, when the green stripe covered almost the entire arena, the butterflies revealed a preferred flight direction towards what now appeared as a dark stripe (n = 18, p = 0.003, MMRT, 8 = 184°, 95% CI [94 - 274°], Fig. 1E, F bottom panel). Overall, we observed a shift in flight directions from flying towards the center of the green stripe, to flying towards the dark stripe center when the stimulus increases in size (Fig. 1G).

### Monarch butterflies rely on a contrast-based orientation strategy

As the butterflies seem to have a preference to fly towards areas of high intensity contrast, i.e., towards the stripe edges, we next focused on studying this in more detail. If we visualized the butterflies’ headings of all experiments in density plots, in almost all cases, the density distribution revealed two peaks, that roughly aligned with the position of the stripe edges (Fig. 2A). We then calculated the absolute angular distance between the butterflies’ heading direction and the centers of the stripes (absolute heading direction) and compared these values to the absolute angular distances between the stripe centers and their respective edges (absolute edge position, Fig. 2B). In this way, we were able to compare the two edges simultaneously, and eliminate potential bimodal heading directions caused by the edges. We observed a clear effect of absolute edge position on absolute heading direction (Median mixed model, p < 0.001, β = 0.60, 95% CI [0.36 – 0.84], Fig. 2B). Thus, the butterflies had a strong preference of flying towards a stripe edge (Fig. 2A, B), irrespective of stripe widths.

To further test whether the individual butterflies switched between edges, or fixated on a single edge, we defined each stripe edge as the stripe width centered at each edge and analyzed the proportion of time spent by each butterfly heading towards each edge (Fig. 2C). Thus, a proportion of 0.5 indicates that the butterflies spent an equal amount of time on both edges during flight, while a proportion that deviates from 0.5 suggests that an individual’s heading choices was biased towards one of the stripes. Here we found that the majority of individuals spend an equal amount of time flying towards each edge, and do not fixate on just a single edge (Fig. 2D, E). The decreased unimodal directedness in the wider stripe phases is then likely explained by the individual butterflies switching between edges, thus resulting in low unimodal directedness (Fig. S1B). These results suggest that Monarch butterflies rely on a simple strategy to keep a constant flight heading with respect to a visual landmark by keeping the landmark’s edge – specifically the stripe – within their frontal visual field, Monarchs enhance the brightness contrast between their left and right eyes, enabling them to maintain a directed course using a contrast-dependent orientation strategy. Thus, the attraction behavior toward a stripe previously reported in Monarch butterflies (Franzke et al., 2022) is not driven by phototactic orientation, but rather by brightness contrasts created by the stripe’s prominent edges. When these edges align along a similar azimuthal direction – as seen with a narrow bright stripe (11°) or a dark stripe (349°) – the butterflies appear to fly toward the center of the stripe, rather than toward its edges.

### Elevation-dependent head-direction system in Monarch butterflies

As opposed to the contrast-dependent strategy observed with a green stripe, where flight directions are biased toward the edges (as shown in Fig. 2), the orientation behavior in response to a small green light spot is fundamentally different. In this case, Monarchs can adopt arbitrary heading directions relative to the stimulus (Franzke et al., 2022). We therefore focused on understanding the conditions under which Monarchs maintain arbitrary flight directions and what exactly triggers the switch from stimulus attraction to menotactic orientation. In a similar fashion to the previous experiments, we presented the butterflies first with a narrow stripe (vertical extent 43°, Fig. 3A). We then decreased the vertical extent of the stripe (22°, 11°), while the stripe width remained the same (11°), until it gradually transformed into a small, green light stimulus (5°) at an elevation of 0° (as measured from the position of the butterflies’ eyes) and finally disappeared (0°). To avoid any history-dependent effects (Beetz and el Jundi, 2023), the order of the vertical stripe extent was presented in a randomized order to the butterflies.

**Fig. 3.**
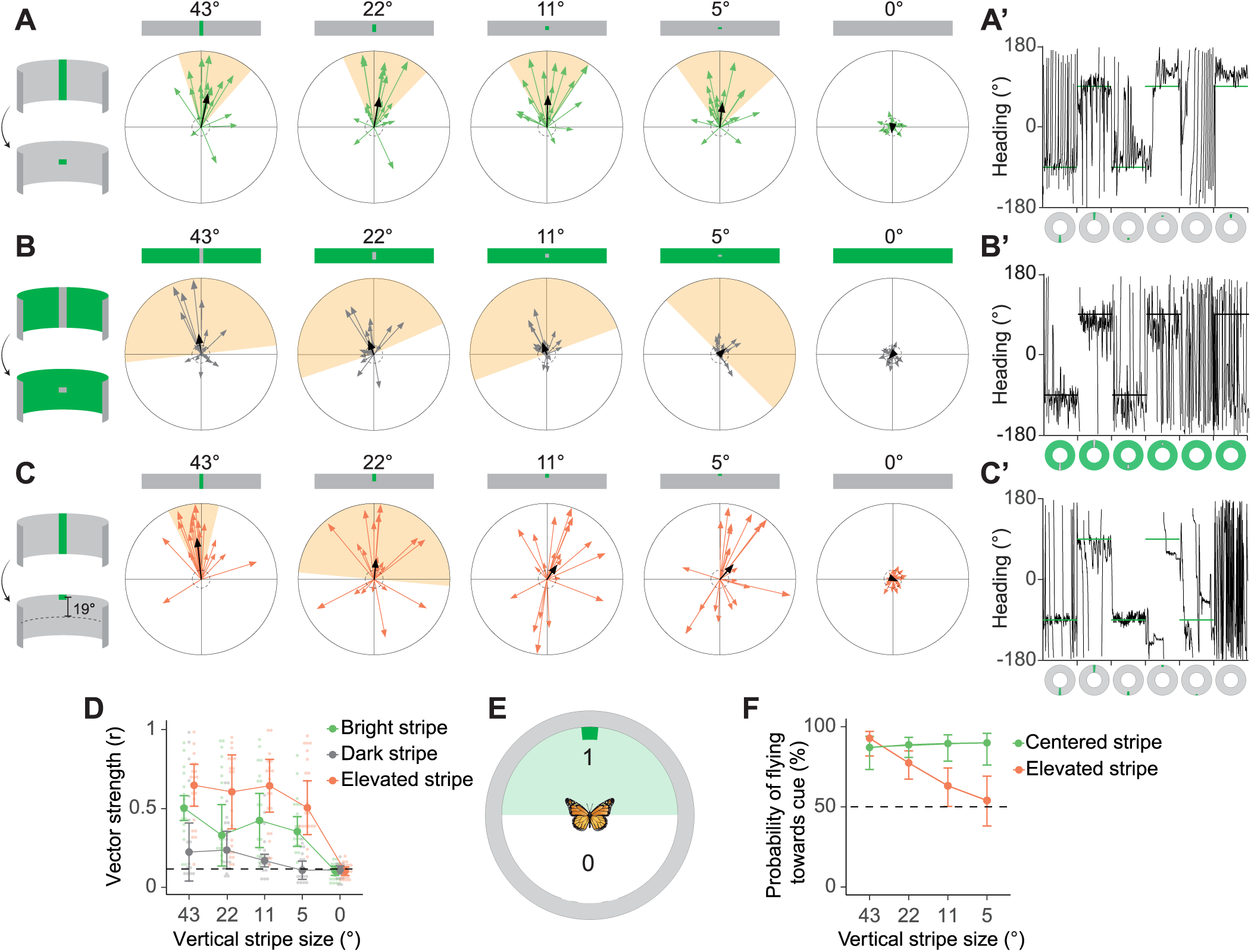
Landmark orientation to a vertical stripe of varying height, contrast, and elevation. (A, B, C) Flight directions and mean vector strengths for individual trials for each experiment (Centered stripe, Dark stripe, Elevated stripe). Thick black arrow shows the population mean vector, supported by the 95% confidence interval of the mean direction (shaded orange area) when the Moore’s modified Rayleigh test for uniformity implies deviations from uniformity (p < 0.05). (A’, B’. C’) Heading directions of exemplary individuals for each experiment. Stripe positions are displayed as a green or dark stripe. (D) The individual vector strengths for each condition, showing the low directedness towards the dark stripe. (E) Illustration of the definition of a binary variable {0,1} used to indicate whether butterflies are flying towards or away from the stripe stimulus. (F) Model predictions from a binomial GLM showing the probability of flying towards the stripe.

As expected, we observed high directedness towards the full stripe stimulus (n = 19, p < 0.001, MMRT, 8 = 12°, 95% CI [342° - 41], Fig. 3A, A’). When we decreased the vertical extent of the stripe, the directedness remained high (Fig. 3A, D), suggesting that these butterflies were well-oriented as long as a visual stimulus was available. Unexpectedly, the butterflies maintained their fixed flight towards the stimulus, irrespective of the vertical stripe extent (Fig. 3 A, A’). Thus, to our surprise, the butterflies did not switch to menotactic orientation (p < 0.001 for all, MMRT). We repeated the same experiment, but, this time, with the inverted visual condition (dark stimulus on a bright background). Again, we observed that the butterflies continued flying towards the stripe, irrespective of stripe height (Fig. 3B, B’). While some individuals exhibited high directedness towards the full dark stripe (Fig. 3B’), the population median 95% confidence interval spans our threshold for directedness (r = 0.22, 95% CI [0.04 – 0.41], n = 18), which decreases as the stripe gets smaller (Fig. 3B, D). Overall, the employed strategy of the butterflies remained the same: They were always attracted towards the cue if provided with a visual stimulus. We therefore conclude that Monarch butterflies do not fly in arbitrary directions with respect to a small light cue presented at an elevation of 0°.

In previous experiments, in which Monarch butterflies exhibit menotactic orientation, the sun stimulus was always presented at a higher elevation (about 23°, Beetz et al., 2023; Franzke et al., 2022). We therefore investigated if elevation might be a crucial filter for the butterflies to interpret a visual stimulus as the sun. Consequently, we now repeated the very same experiment but decreased the vertical extent of the stripe stimulus toward an elevation of about 19° at the smallest vertical extent (Fig. 3C, C’). When we simultaneously decreased the vertical extent of the stripe, and increased the elevation of the cue, the individual directedness remained high, showing an overall well-oriented performance of the butterflies, similar to the conditions with the bright stripe, transferred into a small light cue at 0° elevation (Fig. 3D). However, the butterflies exhibited larger variations in heading directions, resulting in a barely significant group direction with the halved stripe (n = 20, p = 0.034, MMRT, 8 = 5°, 95% CI [275° - 95°], Fig. 3C), and no preferred group direction thereafter (n = 20, p < 0.05, MMRT, Fig. 3C). These results show that the butterflies now performed menotactic orientation to the elevated light stimulus. To compare the two experiments with the different stimulus elevations, we investigated the interaction between experiment type (centered stripe, elevated stripe) and the effect of stimulus size on the probability of flying towards the cue (binomial response {0,1}, Fig. 3E). We observed a significant interaction effect between the experiment type and stimulus size on the probability of flying towards the cue (p = 0.011, binomial GLM), where the centered stripe maintains a flat slope, and the elevated stripe decreases the probability of flying towards the cue, with a confidence interval spanning the expected probability for menotactic behavior at a stripe size of 11° (Fig. 3F). Taken together, Monarch butterflies are attracted by visual cues presented at eye level, and gradually switch to menotactic orientation when a visual cue increases in elevation.

## Discussion

In this study, we investigated the orientation strategies employed by Monarch butterflies when presented with a visual cue whose width, height, brightness, and elevation were systematically varied. Our findings show that the orientation strategy of Monarch butterflies can change instantaneously depending on the perception of a single light cue, a dynamic switch consistent with recent findings (Franzke et al., 2022). Although the perception changes in terms of shape, size, brightness and elevation, these properties may be linked to different ecological interpretations, such as a landing site, or a celestial body.

We observed that Monarch butterflies exhibit attraction to a vertical stripe, a behavior previously documented in Monarchs and other insect species (Franzke et al., 2022; Giraldo et al., 2018; Maimon et al., 2008). Brightness-based attraction may be ecologically advantageous in environments where obscured views of the sun due to cluttered surroundings or dim-light conditions necessitate navigation toward brighter visual areas (Baird and Dacke, 2016). Thus, there is clear ecological motivation for such a sensory-driven mechanism. Notably, as stripe width increased, butterflies continued to approach the stimulus but shifted their focus from the center to the edges of the stripe, a behavior that becomes distinguishable only at wider widths (Fig. 4). This shift indicates a transition from a brightness-based to a contrast-based orientation strategy, wherein butterflies appear to fixate on the boundary between the stripe and its background. This edge-following behavior may enhance orientation precision by leveraging high-contrast visual features, which are often more reliable and stable cues in visual scenes.

**Fig. 4.**
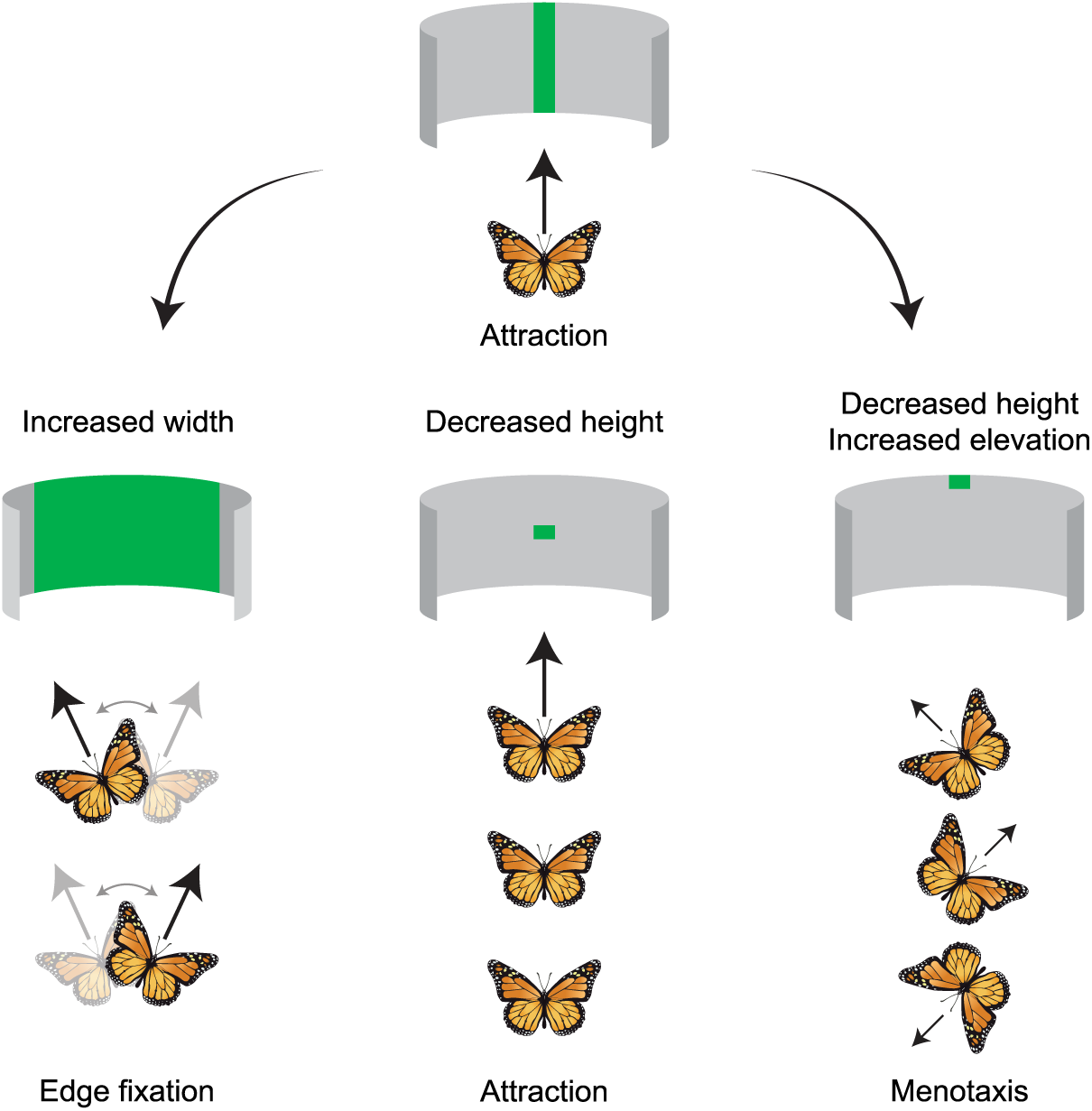
Summarizing behavioral mechanisms in response to different visual stimuli. The butterflies display attraction behavior to a narrow stripe, and a small spot at eye-height. As the stripe width increases, the butterflies fixate on the stimulus edges, switching between them within a flight. When the small spot is elevated above eye-height, the butterflies display menotactic behavior, flying in random directions relative to the stimulus.

### Edge fixation and attraction in Monarch butterflies

Monarch butterflies headed towards the edge of a vertical landmark, the area of maximum brightness contrast. The edge fixation behavior continued irrespective of stimulus width (Fig. 2A) and may explain why attraction behavior is seen towards a dark stripe (Fig. 1E). Further, we observed that individuals switch between the two edges, displaying a low degree of lateralization towards either edge. Edge fixation has also been found in a wide range of other insects ranging from blowflies (Osorio et al., 1990), mealworm beetles (Varjú, 1976), and stick insects (Meschenmoser and Dürr, 2025). Despite this great number of insects relying on the contrast of an edge in laboratory experiments, the ecological relevance for edge fixation is still unclear. Contrast-based attraction could be utilized to maintain a heading direction by using the contrast between distant landmarks and the sky (Möller, 2002), useful for dispersal behavior over longer distances without having to rely on a global compass cue. Interestingly, Monarch butterflies reveal binocular vision in the frontal visual field with most of the photoreceptors detecting visual signals frontally (Kraus et al., 2026). Thus, by keeping the stimulus edge in their binocular field of view they may employ stereopsis to estimate the distance to the stimulus during flight. In combination with optic flow information (Baird et al., 2013), this may contribute to the robustness of several orientation behaviors where insects need to accurately maintain a directed course over short distances, when approaching landing sites and food sources such as stripe-patterned flowers (de Ibarra et al., 2015; Richter et al., 2023) or a mating partner. Alternative to distance measurement via stereopsis, the motivation for an edge-fixation strategy could also be explained from a mechanistic point of view. Thus, it might be computationally a simpler and energetically more effective task for the animal, and thus for the orientation network in the brain, to maintain a heading with high precisions towards an area of high contrast. Thereby, the maintenance of the flight heading could be facilitated by a contrast enhancement mechanism based on lateral inhibition taking place in the retina of the eye as proposed for flies (Osorio et al., 1990). Interestingly, experiments in the fly *Calliphora erythrocephala* revealed that elimination of the frontal visual field, including stereopsis vision, does not affect the visual stripe fixation (Horn and Fischer, 1978). This raises the possibility that other regions of the eye may induce edge fixation. Performing experiments on tethered-flying Monarch butterflies in the future in which different regions of the eyes are painted over may reveal a deeper understanding of the contribution of the visual system to stripe fixation and the underlying mechanistic principles of edge fixation.

When the stripe height was reduced to a small light spot, Monarch butterflies continued to orient toward the stimulus regardless of whether it was a bright spot on a dark background or a dark spot on a bright background (Fig. 3). The latter contrasts with the behavior observed in fruit flies, which exhibit antifixation or avoidance of short, narrow dark stimuli (Maimon et al., 2008). The reduced behavioral performance of the Monarchs in the dark-stimulus condition (Fig. 3B) was expected, as inverting the visual contrast increased the overall brightness intensity with the flight simulator, thereby weakening the salience of the visual cue, similar to what has been observed when presenting a green sun stimulus on a bright background (Franzke et al., 2020). The fact that the animal still maintains their heading to the small light stimulus suggests that stimulus size and shape alone are insufficient to elicit consistent menotactic behavior in Monarch butterflies. However, when the stripe height was reduced at a higher elevation, we observed a progressive increase in the proportion of arbitrary headings, with no preferred group direction at a stimulus size of 11° (Fig. 3C). This adoption of different flight headings is in line with menotactic orientation (Grob et al., 2021), which has previously been documented in Monarchs under similar conditions (Franzke et al., 2022).

### Menotactic orientation to a simulated sun

Like Monarch butterflies, many insects reveal menotactic orientation when presented with a sun stimulus indoors (el Jundi et al., 2014; el Jundi et al., 2016; Mussells Pires et al., 2024). Menotactic orientation is considered to be computationally more sophisticated than attraction behaviors as it requires an animal to compare its desired flight direction with the current flight heading and to constantly compensate for the mismatch between them (Grob et al., 2025; Honkanen et al., 2019). In addition to the sun stimulus, Monarch butterflies can rely on a wide range of visual cues for menotactic orientation, likely also including polarized light (Reppert et al., 2004, but see also Stalleicken et al., 2005). In the Monarch brain, these cues are processed in the central complex (Heinze and Reppert, 2011; Nguyen et al., 2021; Nguyen et al., 2022), the key brain region that governs menotactic orientation based on head-direction (Beetz et al., 2022; Kraus et al., 2026; encode the current flight direction), goal-direction (Beetz et al., 2023; encode the desired flight direction) and steering neurons (Beetz et al., 2023; encode steering commands by comparing the current and the desired direction). In contrast to menotactic orientation, it has been shown that attraction behavior does not appear to require the central complex in insects (Giraldo et al., 2018). As proposed by Franzke et al. (2022), the processing of a bright stripe may bypass the central complex and instead be directly transmitted from the optic lobe to descending motor pathways, triggering brightness-dependent edge and stripe fixation.

### Switching between attraction and menotaxis

Interestingly, our results show a clear shift in orientation strategy: butterflies switch from attraction at low sun stimulus elevations to menotactic orientation at high elevations (Fig. 4). This finding aligns well with previous studies demonstrating that Monarch butterflies adopt menotactic behavior in response to a sun stimulus at higher elevations (Franzke et al., 2022). In nature, Monarch butterflies derive directional information from the sun at higher elevations to disperse or migrate. In a similar way, when our artificial stimulus was presented at an elevation, the stimulus may have similar ecological interpretation as the sun and can be used as a stable cue for menotaxis. Previous studies have also shown that insects associate specific wavelengths to different ecological interpretation, where green light is regarded as a sun stimulus (Brines and Gould, 1979; Edrich et al., 1979; Rossel and Wehner, 1984; el Jundi et al., 2014; el Jundi et al., 2015). Whether light spots of different wavelengths can be interpreted as the sun when presented at an elevation remains to be tested. First data, however, suggest that a UV light stimulus results in a phototactic behavior, irrespective of stimulus elevation (Franzke, el Jundi, unpublished).

One potential neural mechanism underlying this elevation-dependent switch could involve a functional segregation of visual inputs across different regions of the butterfly’s eye. While signals detected by dorsal eye regions, including polarization input from the dorsal rim area (Stalleicken et al., 2006), may contribute to sky compass orientation and promote menotactic behavior, ventral eye regions may process cues related to landing sites or potential mates, thereby driving attraction to visual stimuli. Although direct evidence for such a functional division of visual processing in Monarch butterflies is currently lacking, recent work in *Drosophila* suggests a bias toward greater synaptic input of neurons transferring signals to the central complex in dorsal regions of the medulla (Garner et al., 2024). This may result in a stronger input of signals from the animal’s dorsal field of view to the central-complex network, favoring menotactic orientation at high elevations. Future research, such as detailed mapping of the neural circuits underlying compass orientation in the optic lobe, will be crucial to test this hypothesis and to uncover the precise mechanisms by which visual context shapes navigational decisions in Monarch butterflies.

## Acknowledgments

We thank Keram Pfeiffer and the mechanics workshop of the Biocenter of the University of Wuerzburg for building important pieces of the LED flight simulator. We are also grateful to Robin Grob for discussions and suggestions and for reading the manuscript. In addition, we would like to thank Sergio Siles (butterflyfarm.co.cr) and Marie Gerlinde Blaese for providing us with Monarch butterfly pupae.

## Competing interests

The authors declare no competing or financial interests.

## Author contributions

Study design: FØH, MF, BeJ. Conducting experiments: FØH, MG. Analysis of data: FØH, MG. Interpretation of data: FØH, BeJ. Drafting of the manuscript: FØH, MG, BeJ. Critical review of the manuscript: FØH, BeJ. Acquired Funding: BeJ. All authors approved of the final version of the manuscript.

## Funding

This work was supported by the Norwegian University of Science & Technology, the Emmy Noether program of the Deutsche Forschungsgemeinschaft granted to BeJ (GZ: EL784/1-1) and partly by the DFG under Germany’s Excellence Strategy - EXC 3051/1 “NaviSense” (project number 533653176).

## Data availability

Raw data of all experiments can be downloaded from the website Figshare (https://doi.org/10.6084/m9.figshare.32411424). Analysis scripts can be obtained from the corresponding author upon request.

## Notes

https://doi.org/10.6084/m9.figshare.32411424

### Competing Interest Statement

The authors have declared no competing interest.

