## Supplemental Figure 1 for "Visual cue properties determine innate orientation strategy in Monarch butterflies"

### Supplementary Figure:

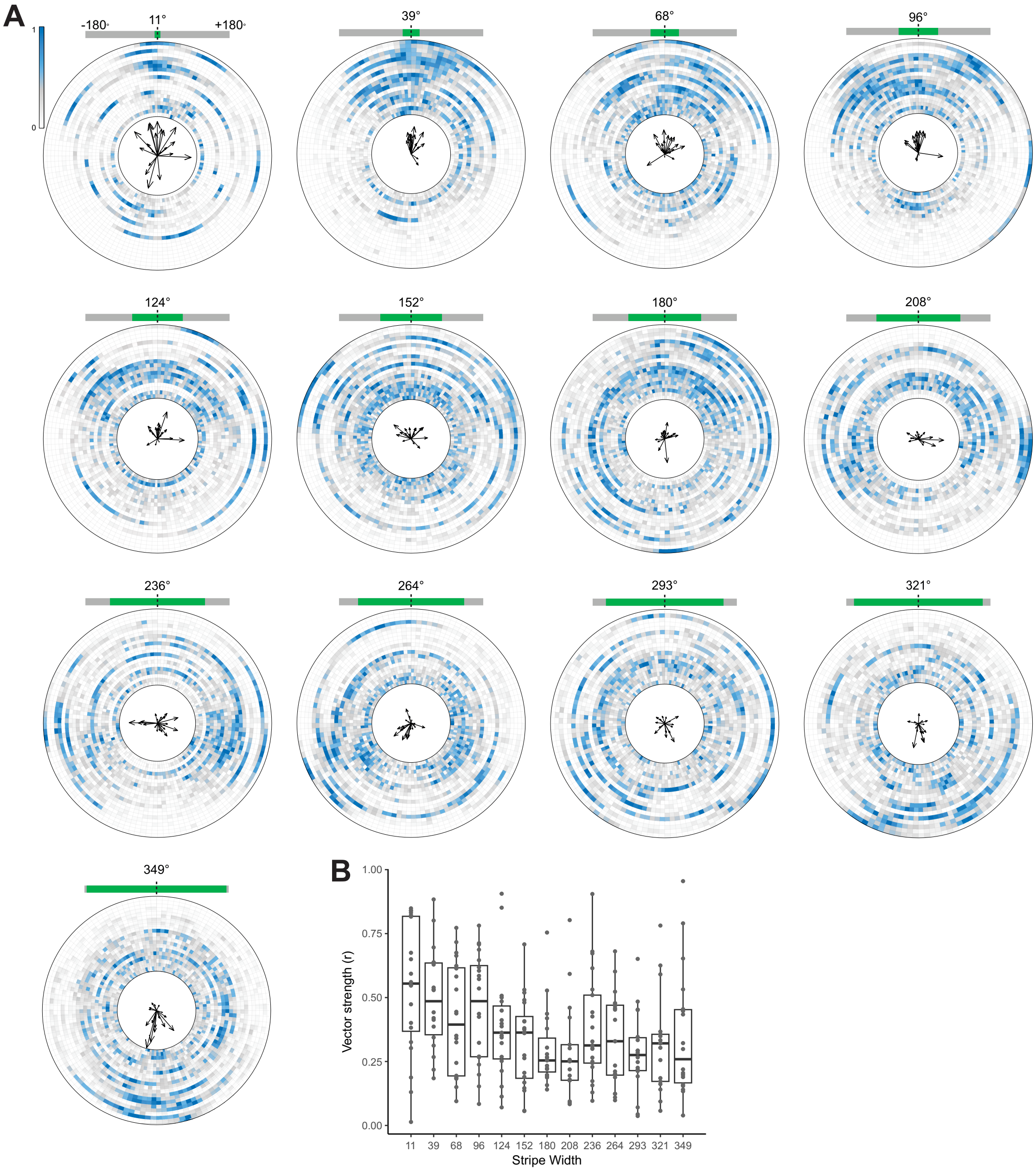


**Fig. S1.** (A) Mean heading directions and vector strengths displayed inside circular heatmaps of raw heading directions for 13 different experimental conditions. The heatmaps display the normalized count of heading directions, with each trial split into separate rings, sorted by increasing vector strength starting from the center. (B) Vector strengths across the different experimental conditions displayed as boxplots, where the horizontal bar within each box displays the population median.
